# RNAi screening of the tyrosine kinome in primary patient samples of acute myeloid leukemia

**DOI:** 10.1101/256040

**Authors:** David K. Edwards, Fatma Eryildiz, Jeffrey W. Tyner

## Abstract

Targeted therapy has proven to be successful in improving outcomes across multiple cancer types. However, many challenges still remain for implementation of this strategy in most patient cohorts, especially with the challenges of identifying the specific mutations or abnormalities in a heterogeneous tumor that are functionally significant. Previously, we developed a functional screening assay, RNAi-assisted protein target identification (RAPID) technology, which evaluates the viability of tumor cells after exposure to siRNAs against members of the tyrosine kinome and NRAS/KRAS. Here, we publish the comprehensive results of this screen for 332 primary AML patient samples. Data from these screening efforts have already helped identify previously unknown therapeutic targets, and will continue to provide insights into better treatment strategies for these patients.

## Introduction

Acute myeloid leukemia (AML), a hematologic malignancy involving the proliferation of immature myeloid cells, is a heterogenous disease.^1,2^ Despite this intrinsic heterogeneity, targeted therapies have proven effective in AML subtypes with particular driver mutations. For example, for AML patients with FLT3 mutations, the kinase inhibitor, midostaurin, was shown to be more effective in combination with chemotherapy than chemotherapy alone.^3^ This finding resulted in midostaurin approval by the US Food and Drug Administration for use in newly diagnosed AML patients. There are numerous other efforts to implement this approach across different molecular subtypes, with varying degrees of success.^4^

One limitation to the implementation of this targeted therapy approach is in determining the molecular targets in AML that are functionally significant. To overcome this limitation, we have developed a method of screening primary AML patient cells, which we termed RNAi-assisted protein target identification (RAPID).^5,6^ Partial results from this screening technique have been published elsewhere,^7-14^ although the entire dataset has never been released.

Here, we are publishing the complete dataset containing the results of the RAPID screen performed on 332 primary AML patient samples. The significance of many of these kinases in the pathobiology of AML are currently being studied in our group and elsewhere.

## Methods

Detailed methods for the RNAi-assisted protein target identification (RAPID) functional assay, including the electroporation conditions and the siRNA sequences, have been published elsewhere.^5,6^ Briefly, peripheral blood, bone marrow aspirates, or leukapheresis samples were obtained from AML patients by informed consent according to a protocol approved by the Oregon Health & Science University (OHSU) Institutional Review Board. The peripheral blood mononuclear cells (PBMCs) were isolated from these samples using a Ficoll density gradient. These cells were added to a 96-well plate containing 91 arrayed siRNA pools that collectively target the tyrosine kinome, as well as siRNA pools targeting NRAS and KRAS. Following electroporation, the cells were transferred to basic cell culture media and incubated for 96 hours, at which time the cell viability was determined by a tetrazolium-based MTS assay.

We analyzed the data as reported previously.^5,6^ Briefly, for each sample, we classified “hits”, or siRNAs that had a significant impact on cell viability, as those whose viability was less than two standard deviations below the mean viability for all siRNAs on a given patient sample run. We performed unsupervised hierarchical clustering on the “hits” for each patient sample using Ward’s method for clustering (the “hclust” function). The data was collected, analyzed, and visualized in R.

## Results and Discussion

In order to understand the results from our RNAi-assisted protein target identification (RAPID) functional assay, we graphed the frequency of “hits” that were observed in our 332 primary AML patient sample cohort (**Figure 1**).

**Figure 1.**
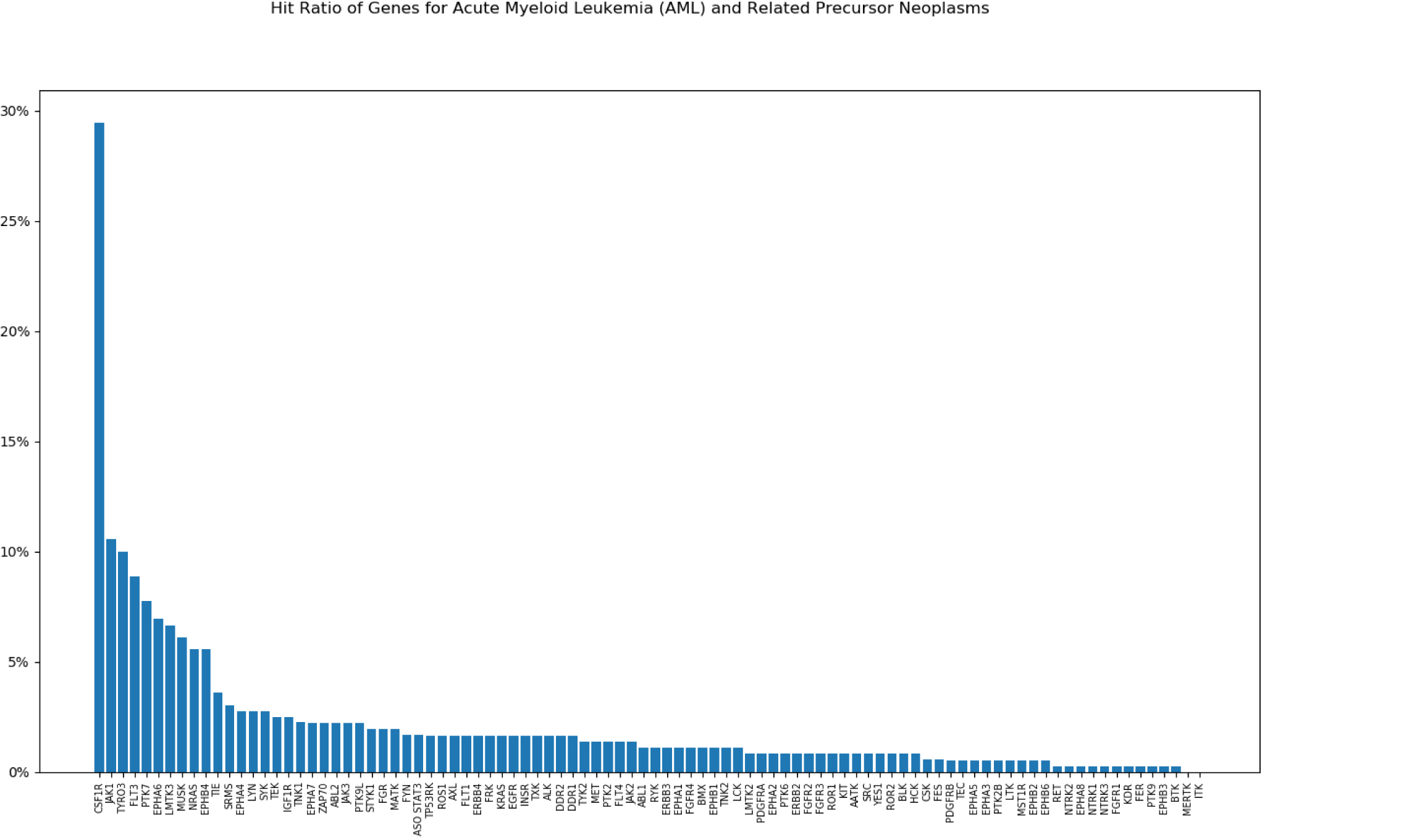
<u>Frequency of kinase “hits” in primary AML patient samples.</u>. Kinase targets and NRAS/KRAS screened using the RAPID assay are displayed on the x-axis and percentage of patients that showed significant sensitivity to the corresponding target siRNA exposure on the y-axis.

The most significant “hit” in our screen was against CSF1R (colony stimulating factor 1 receptor), a class-III receptor tyrosine kinase that is expressed on monocytes and terminally differentiated macrophages. The large-scale sequencing effort of 200 *de novo* AML patients undertaken by The Cancer Genome Atlas analysis did not identify any mutations in CSF1R, nor is it significantly overexpressed.^1^ (It should be noted that there are two cases of CSF1R mutations/translocations identified in AML cell lines— specifically, the Y571D mutation in GDM-1, which results in constitutively active CSF1R,^15^ and the t(3;5)(p21;q33) *RBM6*-*CSF1R* translocation in MKPL-1, which renders it sensitive to CSF1R-targeting small-molecule inhibitors.^16^ However, these mutations have not been observed in any additional patient samples.) Our lab recently presented data to suggest that the significance of CSF1R in AML is related to the tumor microenvironment,^17^ and we are working further to uncover more information about the mechanism of action.

The second most significant “hit” was against JAK1 (Janus kinase 1), a non-receptor tyrosine kinase involved with the activity of numerous cytokine signaling pathways. The significance of JAK1 mutations in AML is poorly understood. In a screen of the entire JAK1 coding region in AML patients, 2 out of 94 patients were found with JAK1 mutations.^18^ Although overexpression of these mutants did not transform Ba/F3 cells, they did show activation of downstream STAT1 and other signaling pathways.^18^ In addition, small-molecule inhibitors of JAK1 (and its related family member JAK2) are currently being investigated in relapsed or refractory AML patients.^19,20^ The role of JAK1 activation in the context of our patient sample cohort, however, is not fully understood.

The third most significant “hit” was against TYRO3 (TYRO3 protein tyrosine kinase), a member of the TAM family of receptor tyrosine kinases with a conserved extracellular structure and kinase domain sequence. Not much is known about the significance of TYRO3 in AML, although it has been shown to be upregulated in AML^21^ and targeting similar family members is an active area of investigation in leukemia.^22^ We recently presented some data on the significance of TYRO3 in AML,^23^ although we are conducting more research to uncover its full significance.

Finally, the fourth most significant “hit”, was against FLT3 (fms related tyrosine kinase 3), a class-III receptor tyrosine kinase whose involvement in AML has been well-studied. FLT3 is one of the most commonly mutated proteins in AML,^1,2^ appearing either as an internal tandem duplication (FLT3-ITD) or a mutation in the tyrosine kinase domain (FLT3-TKD). This result is not unexpected, therefore, and serves as an indication that the results we observe in our patient sample screening conform with existing knowledge of AML pathobiology.

Finally, to identify any patterns of co-occurrence between siRNA hits among the patient samples, we performed unsupervised hierarchical clustering on the dataset (**Figure 2**). Overall, there were no significant clusters observed among the siRNAs, suggesting that each tyrosine kinase functioned independently of the activation of other tyrosine kinases. This result is not surprising, considering that there is a long-standing hypothesis that two classes of mutations are required to be present for leukemogenesis: disease-initiating mutations that block cell differentiation, and disease-progressing mutations that enhance cell proliferation and survival.^24^ Many of the disease-progressing mutations in AML occur in tyrosine kinases, specifically activating mutations in FLT3, KIT, and NRAS,^25^ so the activation of two kinase pathways could be functionally redundant for the development of leukemia.

**Figure 2.**
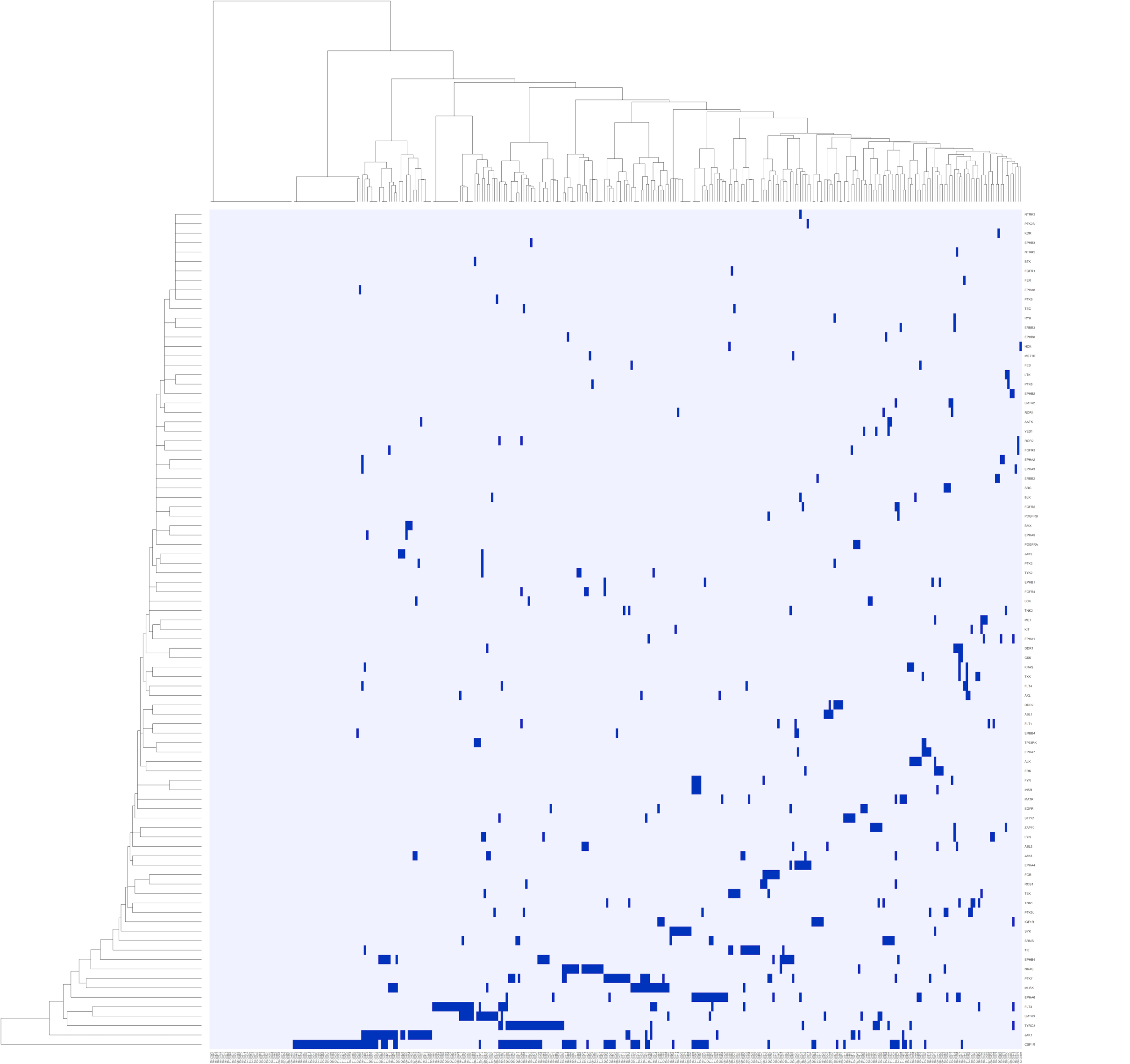
<u>Heat map of kinase “hits” in primary AML patient samples.</u>. Patient specimen IDs are displayed on the x-axis and tyrosine kinases and NRAS/KRAS from the RAPID screening panel are displayed on the y-axis. siRNA “hits” are colored dark blue.

## Figure Legends

<u>Supplemental Table 1. Complete dataset of RAPID screening results in primary AML patient samples.</u> White blood cells from 332 AML patients were processed and the sensitivity for each individual siRNA knockdown were measured as previously described. Data points below two standard deviations of the mean cell viability (at *P*<0.05) were considered “hits”. Data shown here are the viability scores for each kinase from every plate per patient.

